# Molecular pixelation of the CAR T cell surface proteome

**DOI:** 10.64898/2026.01.30.702970

**Authors:** Anthony Cesnik, Oliver Takacsi-Nagy, Trang Le, Theodore L. Roth, Ansuman T. Satpathy, Emma Lundberg

**Affiliations:** Bioengineering Department, Stanford University, Stanford, CA, USA; Biohub, Chicago, IL, USA; Department of Pathology, Stanford University, Stanford, CA, USA; Program in Immunology, Stanford University, Stanford, CA, USA; Science for Life Laboratory and KTH Royal Institute of Technology, Stockholm, Sweden

## Abstract

Immunotherapies using CAR T cells are revolutionizing B-cell acute lymphoblastic leukemia treatments. However, the majority of patients remain unresponsive, and chronic stimulation of T cells is a common contributor that reduces effector function and persistence. We apply Molecular Pixelation, a recently developed single-cell technology for characterizing cellular surface proteomes, to determine characteristic topological surface-based proteomic signatures of CAR T cell exhaustion. We analyze 76 surface proteins on 8504 CAR T cells at a single-cell level, collected from three donors and either stimulated once or repeatedly, six times over two weeks. The abundances, polarizations, and colocalizations of surface proteins can each distinguish CAR T cells that were stimulated acutely or chronically, and all but one marker with polarization changes increased in polarization. These data also reveal disrupted adhesion signatures of protein colocalization in the peripheral supramolecular activation complex (pSMAC) and increased CD37/CD82 colocalization after chronic stimulation. These Molecular Pixelation results convey new spatial signatures for proteomic polarization and colocalization on the cell surface that represent new cell-state axes for immunology and systems biology.

## Introduction

Immunotherapies using chimeric antigen receptor (CAR) T cells have cured refractory leukemias in a subset of patients^1^. However, the majority of patients remain unresponsive to this treatment across broader indications. While there are many mechanisms for this resistance, the intrinsic dysfunction or exhaustion of CAR T cells is a common contributor that diminishes effector function and persistence^2^. CAR T cell exhaustion has been profiled using sequencing and proteomics tools that have revealed valuable insights about this cellular phenotype, but the changes in spatial distribution of proteins on the cell surface remain invisible to these methods. Molecular Pixelation is a recently developed optics-free method to map the topology of protein distributions on the cell surface that was demonstrated in chemokine-stimulated T cells^3^. Here, we use this method to investigate the spatial signatures for proteomic polarization and colocalization on the cell surface in T cell exhaustion.

Chronic stimulation with a tumor antigen is characterized by the loss of effector function, diminished proliferative capacity, persistent TCR signaling, increased levels and coexpression of inhibitory receptors, transcriptional and epigenetic reprogramming, and metabolic dysregulation^4,5^. While inhibitory receptors like PD-1 and TIM-3 serve as important extracellular markers of exhaustion, they lack resolution for earlier exhausted states. Therefore, intracellular markers with a clearer connection to the mechanistic development of chronically stimulated phenotypes have served to identify chronically stimulated T cells, namely transcription factors like TOX and EOMES.

Studies of chronically stimulated CAR T cell phenotypes have mainly been profiled with bulk and single-cell RNA sequencing^2^ and proteomic tools including CyTOF^6^, flow cytometry, and immunohistochemistry (IHC). While IHC can provide spatial information for the dissection of CAR T cell phenotypes *in situ*, the spatial distribution of proteins on the cell surface is currently invisible to current methods. In the present study, we analyze the abundance, polarization, and colocalization of the surface proteome using a recently developed antibody-based spatial proteomics technique named Molecular Pixelation^3^. We use these results to determine how the surface proteome of CAR T cells changes during chronic stimulation. We find that there is a broad increase in polarization of all markers, or in other words the receptors form clusters on the surface of the cell after chronic stimulation. Furthermore, there are characteristic changes in colocalization of surface markers that provide a clearer molecular picture of the cell surface during chronic stimulation.

### Molecular pixelation of the CAR T cell surface proteome

Molecular Pixelation^3^ is an optics-free spatial proteomics technique that maps the topology of protein localizations across the surface of individual cells using DNA sequencing instead of microscopy (Figure 1). After fixation, cells are stained with a panel of DNA-conjugated antibodies (against 76 target proteins and 4 controls in this study). The DNA oligonucleotides encode which protein the antibody targets, and a unique molecular identifier (UMI) to allow unique binding events to be counted. Two sequential rounds of DNA “pixels,” nanometer-scale single-stranded DNA products, are generated by rolling circle amplification. In the first round, DNA pixels hybridize to a universal sequence on nearby DNA-conjugated antibodies, and then a gap-fill ligation reaction copies the pixel’s unique pixel identifier (UPI-A) onto the oligonucleotide conjugated to the antibody. Antibodies that are close enough to the same pixel get tagged as belonging to the same local neighborhood. After the first pixel set is enzymatically removed, a second pixel repeats the process, adding a second neighborhood tag (UPI-B). After PCR and sequencing, each read contains the UMI, protein identity, and neighborhood memberships on the cell surface. Computationally, the link between the two neighborhood memberships is treated as an edge labeled with a protein identity, and the pattern of partially overlapping pixel memberships is used to create a graph informing us of the relative proximity of proteins on the cell surface. The Leiden clustering algorithm is applied to this graph to separate individual cell surface topologies without imaging or isolating cells into droplets during the assay (Figure 2A).

**Figure 1.**
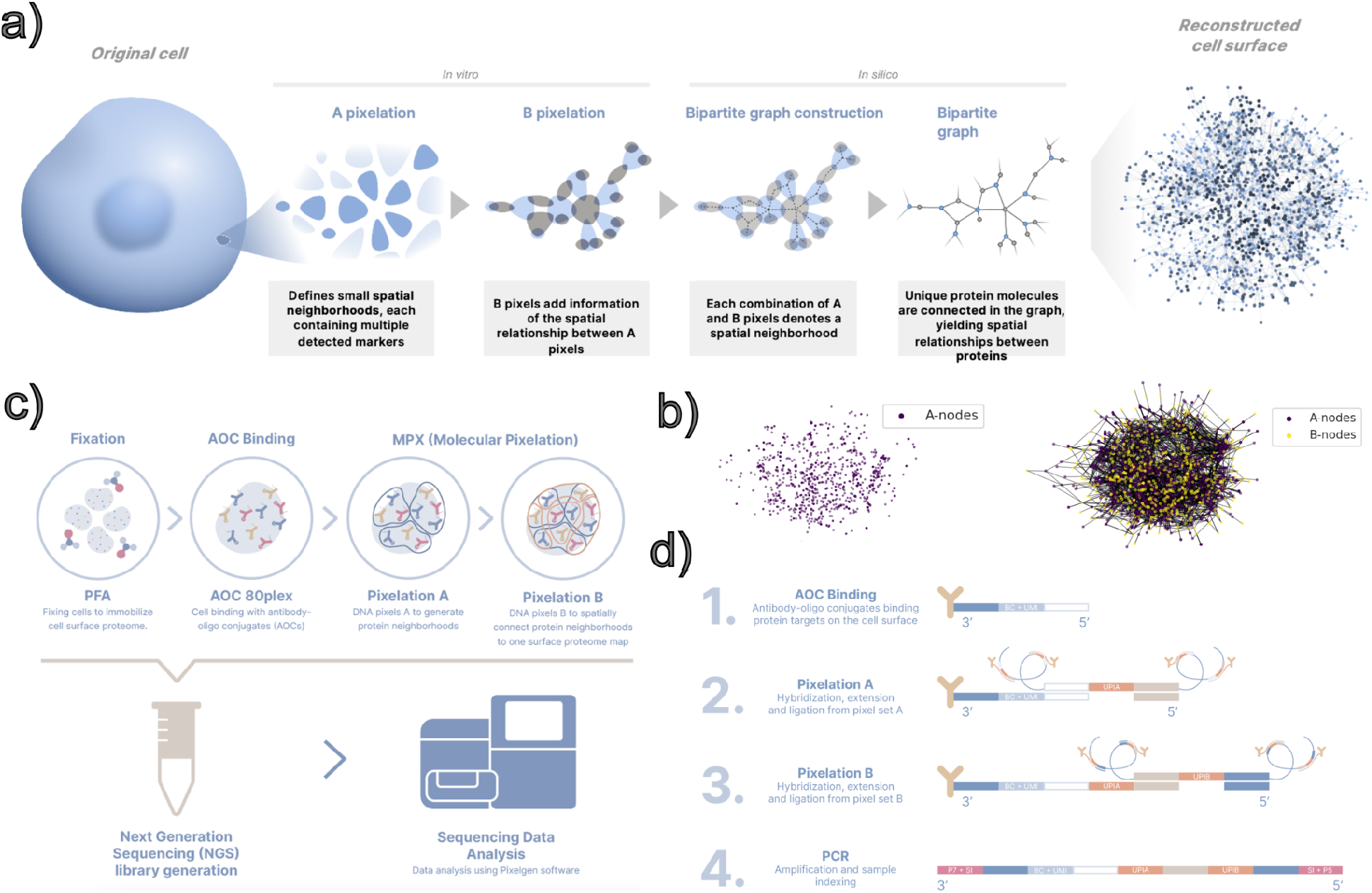
Schematic of Molecular Pixelation. a-b) Two rounds of pixelation are used to map neighborhoods of protein proximities on the cell surface, which are reconstructed as a bipartite graph representing these spatial relationships. c-d) Molecular Pixelation is an optics-free method using antibody-oligonucleotide conjugates (AOC). Two rounds of extension and ligation to these oligonucleotides create pixels that are sequenced and used to reconstruct the cell surface protein proximities.

**Figure 2.**
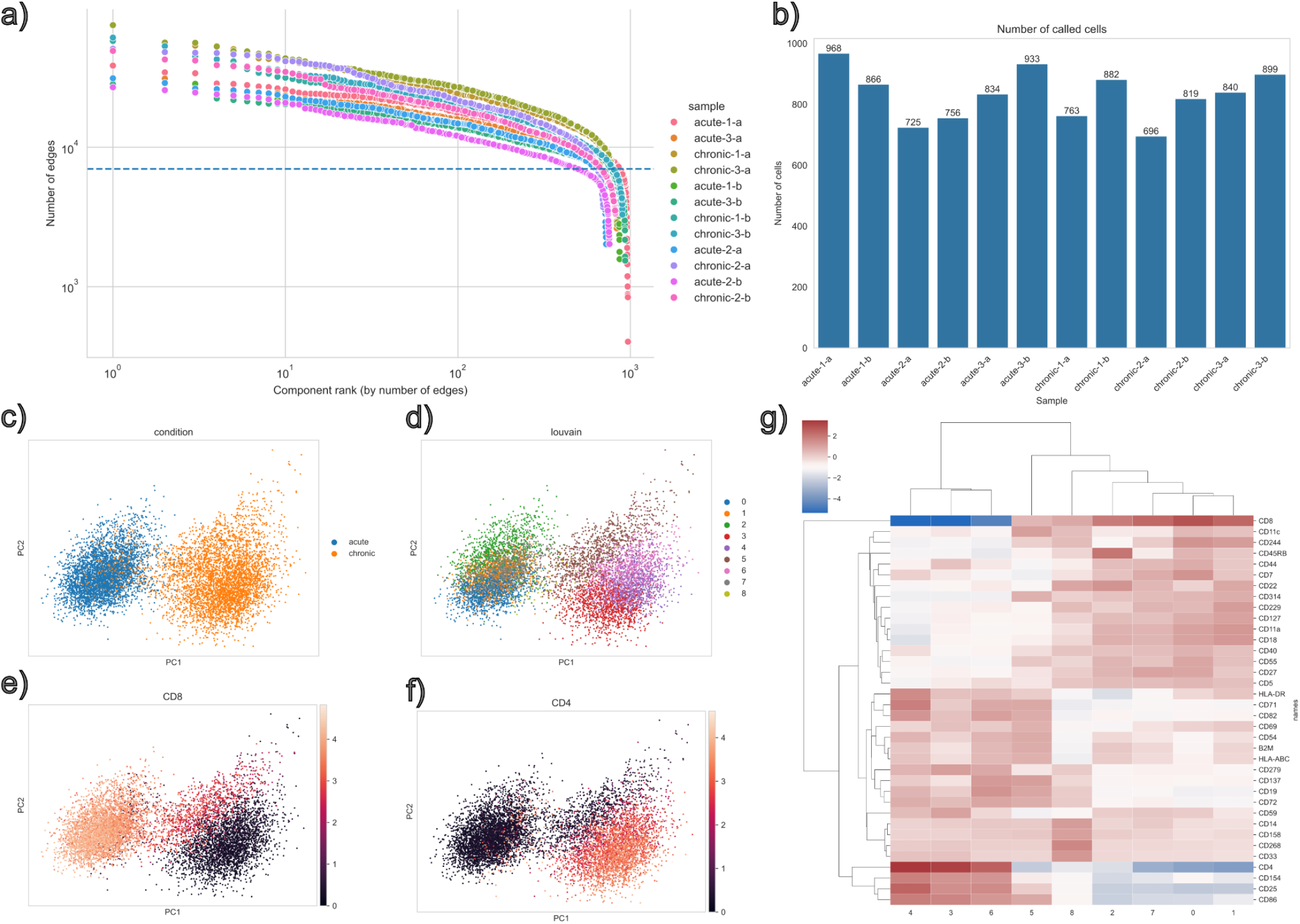
a) Cells are called^3^ from the Molecular Pixelation data based on the count of edges within components (components with >7,000 edges were considered cells). b) Several hundred cells were analyzed in each sample. c-f) Dimensionality reduction using UMAP, overlaid with the condition, either acute and chronic stimulation (c), clusters determined using louvain clustering (d), expression of CD8 (e), and the expression of CD4 (f). g) Significant differential expression was observed across the clusters.

We used this method to analyze 8504 CAR T cells at a single-cell level (Figure 2B). These cells were collected from an *in vitro* model for chronic antigen stimulation^7–9^ that drives T cells towards a dysfunctional exhaustion-like state that is observed with persistent antigen exposure in cancer. Primary human CD3+ T cells were isolated from PBMCs from 3 donors, activated with CD3/CD28 agonism, and expanded in IL-2-supplemented T cell medium. The cells were then gene-edited 48 hours after activation with a template encoding CD19-28ζ CAR (CD19-targeting receptor with CD28 costimulation and CD3ζ signaling) and a truncated NGFR reporter (tNGFR, a biologically inert surface marker for enriching CAR-expressing cells). The edited CAR T cells were expanded and challenged with CD19-expressing Nalm6 leukemia target cells. Stimulation began six days after editing, or eight days after the initial activation, by co-culturing CAR T cells with Nalm6 targets. These cultures were then sampled and re-seeded every 2-3 days based on flow cytometry, using tNGFR to quantify CAR T cells and CD19 to quantify Nalm6, to maintain the effector-cell to target-cell ratio across rounds. “Acute” stimulation in this study refers to cells harvested after one stimulation round, whereas “chronic” stimulation refers to cells stimulated six times over two weeks of repeated re-challenge with fresh target cells. At these experimental endpoints, live tNGFR+ CAR T cells were enriched by flow cytometry for analysis with Molecular Pixelation.

The marker expression results from the Molecular Pixelation analysis separated acutely and chronically stimulated cell states (Figure 2C). When clustering the expression, we found CD4+ and CD8+ CAR T cells in both acutely stimulated and chronically stimulated groups (Figure 2D-F). Chronically stimulated CD8+ CAR T cells (Figure 2D-E, cluster 5) are of particular interest to the study, since they are the cytotoxic cells that attack the target cells. When inspecting markers distinguishing chronically stimulated CD8+ cells from acutely stimulated ones (Figure 2G, cluster 5; Table S2), we find lower abundances of CD44, CD59, CD7 (p = 2.2e-112, 1.3e-159, 1.9E-243), and elevated levels of CD137 and CD11c (p = 2.8e-47, 1.4e-187). These results and those in following sections exclude effects that are likely confounded by trogocytosis where receptors from the Nalm6 cells used to stimulate the CAR T cells are pulled off by the T cells are excluded (see Methods).

### Protein polarization broadly increases after chronic stimulation

A total of 35 of the 76 proteins (46.1%) exhibited significant changes in polarization when all CAR T cells are evaluated as a whole (Figure 3A). Interestingly, all but one of these 35 proteins exhibited increases in polarization after chronic stimulation. This broad increase in polarization may result from diminished active processes required to delocalize proteins and remix the membrane, such as inhibited actin remodeling^10^ and decreased ATP supply due to mitochondrial dysfunction^11^.

**Figure 3.**
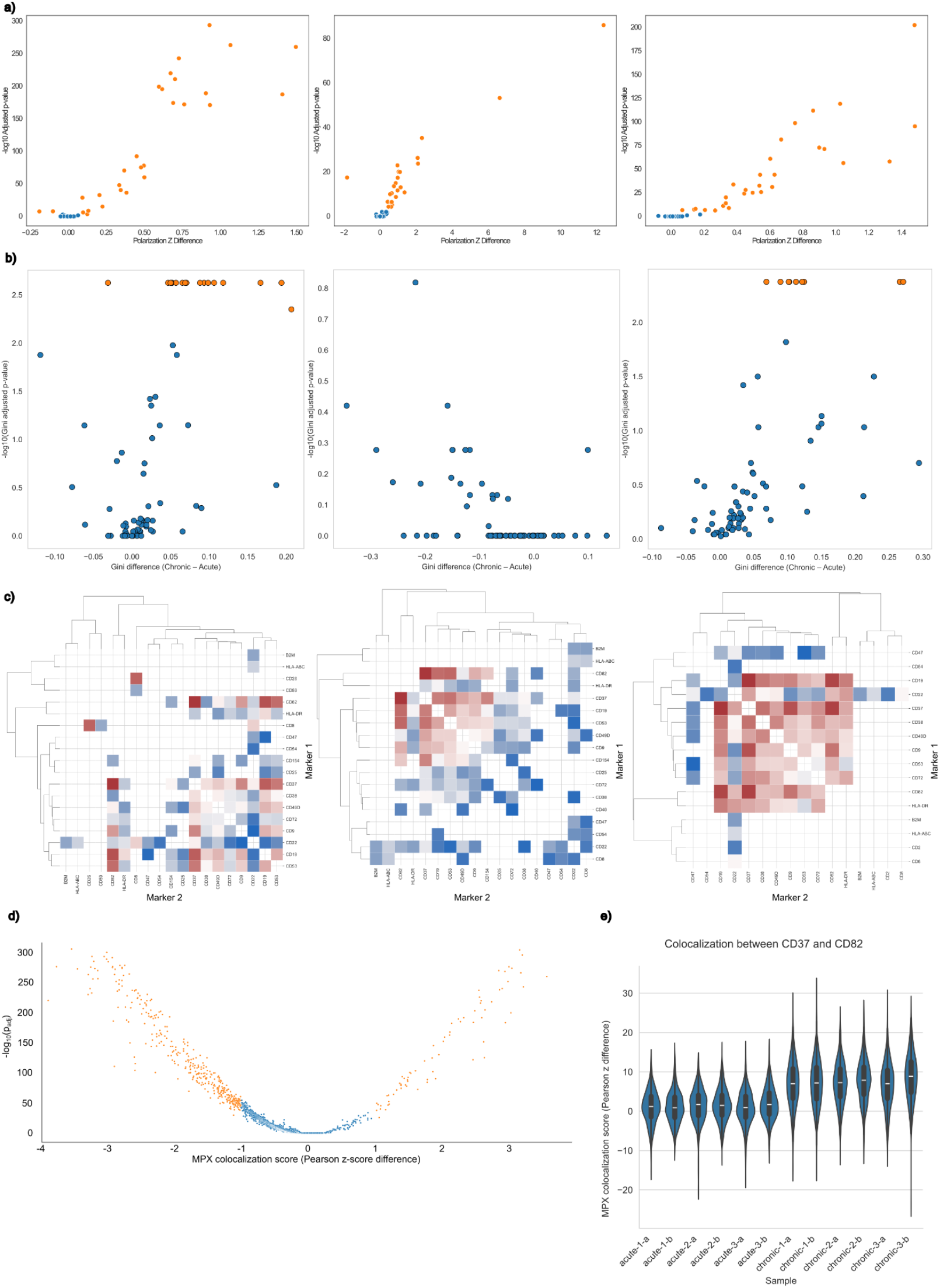
a) Differential polarity of all CAR T cells (left), CD4+ CAR T cells (middle), and CD8+ CAR T cells (right), where orange represents significant polarization changes during chronic stimulation relative to acute stimulation. b) Variability of single-cell polarization of all CAR T cells (left), CD4+ CAR T cells (middle), and CD8+ CAR T cells (right), where orange represents significant changes in the single-cell variability of polarization (gini index) during chronic stimulation relative to acute stimulation. c) Clustermap of significant differential colocalization of all CAR T cells (left), CD4+ CAR T cells (middle), and CD8+ CAR T cells (right), where reds correspond to increases in colocalization during chronic stimulation relative to acute stimulation and blues correspond to decreases. d) Volcano plot showing the 561 protein pairs (19.7% of all 2850 combinations total) in orange that have significant changes in colocalization; non-significant pairs are shown in blue. e) Distributions of CD37/CD82 colocalizations shown per sample and replicate display the reproducibility of the observation of increased colocalization of these markers across samples.

Broader cell type-specific changes were observed in marker polarization than from abundance alone. When CD8+ CAR T cells were evaluated independently, 31 of the 76 proteins (40.8%) displayed these significant changes in polarization, all of which were increases in polarization during chronic stimulation. Similarly, for CD4+ CAR T cells, 24 of the 76 proteins (31.6%) had significant changes in polarization during chronic stimulation. Unique displays of polarization changes were observed for both CD8+ and CD4+ cells, including 7 of the 31 markers for CD8+ cells (22.6%; CD11a, CD137, CD18, CD3E, CD48, CD50, and CD47) and 1 of the 24 markers for CD4+ cells (4.2%, CD40). Of these, only CD47 is overrepresented in NALM6 B cells and is potentially the result of trogocytosis.

### Protein polarization and colocalization of protein pairs distinguishes chronic and acute stimulation

Cellular rewiring is a growing area of study, with reports of over 50% of protein-protein interactions differing between cell types^12^. The changes over the course of cell perturbations are still largely unexplored, and the Molecular Pixelation data allows us to take a look at changes in proximity over chronic CAR T cell stimulation. Significant colocalization changes were observed for 561 protein pairs (19.7% of all 2850 combinations; Figure 3D). One particular colocalization event that stood out was CD37/CD82, which increased significantly after chronic stimulation (Figure 3E, Figure 4; adjusted p-value < 1e-250).

**Figure 4.**
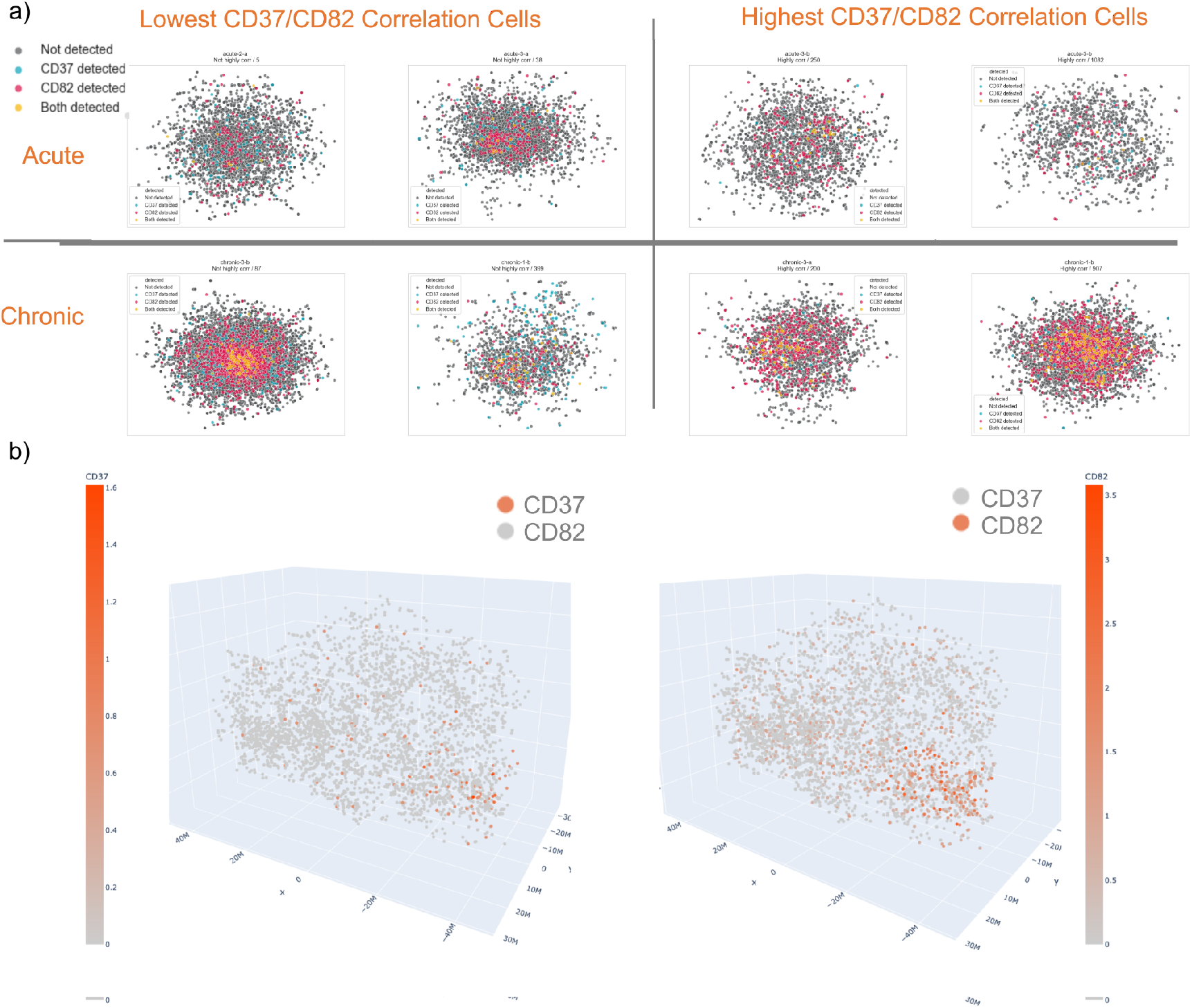
Comparisons of the colocalization of CD37/CD82, where each point is one pixel from Molecular Pixelation. a) Displayed in both acutely and chronically stimulated CAR T cells for two example cells chosen to show both high and low correlation between the markers within the two conditions. These illustrate the stronger correlation of colocalization in chronic stimulation. b) Displayed side-by-side in a 3D rendering of the same chronically stimulated cell, displaying the strong colocalization of CD37 and CD82.

As for polarization, many more cell type-specific colocalization changes were observed than marker abundance alone. For CD8+ CAR T cells, 146 of the 450 protein pairs (5.1% and 15.8% of all 2850 combinations; Figure 3C) were uniquely colocalized and not represented in CD4+ CAR T cells, which had 103 of 561 protein pairs that were had uniquely significant colocalizations (3.6% and 19.7% of all 2850 combinations; Figure 3C). The CD37/CD82 colocalization that is highlighted had significant increases for both types of CAR T cells.

### Destabilization of the immune synapse over chronic stimulation

When we inspect the changes of colocalization in context of the immune synapse, we observe broad delocalization of related markers (Figure 5). We observed delocalization of proteins at the peripheral supramolecular activation cluster (pSMAC), with CD54 and CD2 showing decreased colocalization with CD11a. Thus, the global remodeling of colocalization may impact the immune function of the cells during chronic stimulation and may underlie that phenotype.

**Figure 5.**
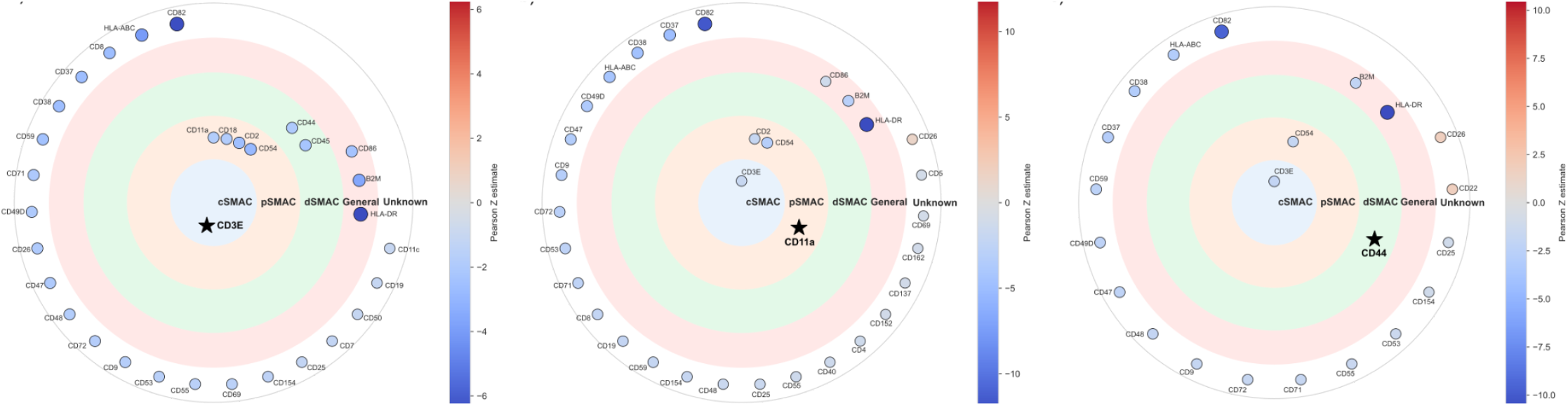
Significant changes in colocalization within the immune synapse of CD8+ CAR T cells, relative to one chosen marker per plot. These include the a) cSMAC represented by CD3E, b) pSMAC represented by CD11a, and c) dSMAC represented by CD44. Protein markers with general and unknown association with the immune synapse are also labeled. These results show destabilization of the immune synapse with delocalization of related markers, particularly with CD54 and CD2 showing decreased colocalization with CD11a in the pSMAC.

The variability of protein expression and localization is known to be widespread between individual cells and impacts phenotypes such as metabolism^13,14^. We observe a general increase in the variability of polarization and over chronic stimulation (Figure 3B) that are consistent across donors (Supp Fig 5), with markers such as CD137, CD59, CD7, and CD82 showing increases in the variability of polarization (adjusted p-values = 0.0042 for each marker, permutation analysis). The increased variability of CD82 polarization in single-cells is intriguing in light of the strong delocalization with the immune synapse and strong increase in polarization with CD37 over the process of chronic stimulation. These increases in variability during chronic stimulation may be another indication of the destabilization of the immune synapse.

### Limitations of study and perspectives for surface topology cell-state measurements

This study used three donors to interrogate these differences. However, this study does have several limitations. There is not a time course to observe the process of chronic stimulation evolve, and the resolution of the cSMAC is limited with only CD3E representing this structure. Future investigations with other components of this central structure, as well as expanding the panel of markers to characterize the localization of the CAR construct itself, will help to give more resolution to these changes.

Molecular Pixelation is a new spatial proteomics technique for the discovery of proteomic polarization and colocalization on the cell surface that represent a new cell-state axis for systems biology and immunology. These observations of changes in protein neighborhoods could also be considered a complex phenotypic marker that could be read out with targeted proximity assays in future studies or even clinical settings. In this study, we observed global restructuring of the cell surface proteome during chronic antigen presentation to CAR T cells, including disruption of components involved in adhesion at the pSMAC of the immune synapse. Other markers, including CD82, which is also involved in adhesion^15^, showed significant changes in polarization, colocalization, and delocalization with the immune synapse in chronic stimulation. These changes may represent a coordinated inhibitory topology, or they may result from diminished active processes required to delocalize proteins and remix the membrane. Overall, these observations serve as candidates for future investigation of CAR T cell dysregulation.

T cell exhaustion plays a role in determining the success of immunotherapies such as checkpoint inhibitors (e.g., PD-1 blockade)^16^. Better understanding how persistent stimulation of T cells induces exhaustion, and how checkpoint blockades modulate these exhausted programs, could help guide therapeutic strategies and efficacy. The study also points to important applications in studying cell signaling. Here, the CAR T cells are responding to the cancer cells used to stimulate their activation, but the results of this study also underscore the importance of understanding the surface proteome dynamics of many cell signaling events. Ultimately, single-cell surface protein proximity measurements of critical interaction surfaces may help us to model cell function for T cells and other cell types by including this type of data in efforts to build foundation models to understand the immune system^17^ and cell biology^18^.

## Materials and Methods

### CAR T cell generation

CAR T cells were generated and assayed as previously described^9^. Briefly, PBMCs were isolated from leukoreduction chambers from three healthy human donors (Table S1) and subsequently CD3+ T cells were isolated by negative selection (StemCell). CD3+ T cells were activated using CD3/28 agonism (Dynabeads - Thermo FIsher) for 48 hours at 1e6 cells/mL of T cell medium (TCM): X-VIVO 15 medium (Lonza) supplemented with 5% FBS and 50 IU/mL human IL-2. Following activation, Dynabeads were magnetically removed and T cells were centrifuged for 10 minutes at 90xg and then resuspended in P3 Buffer (Lonza) at a density of 1-2e6/20 uL. T cells were mixed with Cas9 ribonucleoprotein and a plasmid homology-directed repair template and aliquoted into a 96-well Nucleocuvette Plate (Lonza) for electroporation by pulse code EO-151 using a Gen2 Lonza 4D instrument with a 96-well plate attachment. T cells were immediately resuspended with 80 ul TCM, incubated at 37°C for 15 minutes and then transferred to a standard 96-well plate with a total volume of 300 ul TCM. T cells were maintained at a density of 0.5e6-1e6/ml and TCM refreshed every two to three days.

The plasmid homology-directed repair template encodes a multicistronic cassette with a tNGFR gene and a CD19-28ζ CAR, flanked by sequence homology to the TRAC locus, enabling in-frame integration. Edited cells were enumerated by flow cytometry using an antibody against tNGFR and counting beads (CountBright Plus Absolute Counting Beads - Thermo Fisher).

At six days following editing, CAR T cells were co-cultured in TCM with Nalm6 cells at an effector to target ratio of one to eight. Two days later, half of the culture was restimulated with Nalm6 cells at the same E to T ratio (“Chronic”), while the other was maintained in TCM for the remainder of the assay (“Acute”). The chronic culture was restimulated 4 additional times for six total stimulations over the course of two weeks. At the end of the assay, live and tNGFR+ cells were enriched from both culture conditions by FACS for downstream analysis.

### Molecular Pixelation

Cells were processed using the Pixelgen Single Cell Spatial Proteomics Kit (Pixelgen Technologies, PXGIMM001) according to the manufacturer’s protocol and recommended reagents. In short, cells were thawed, washed, counted and fixed using 1% PFA (Electron Microscopy Sciences, 15710). After blocking, the samples were split into duplicates and stained with an 80-plex oligo-conjugated antibody panel, followed by stabilization with a secondary antibody. From each sample replicate, 20,000 stained cells were then subjected to Molecular Pixelation. Molecular Pixels A were added, followed by a gap-fill reaction and subsequent Pixel A removal. Then, Pixels B were added, followed by a second gap-fill reaction. The cells were counted and 1000 cells from each sample replicate were subjected to exonuclease treatment and indexing PCR. The PCR products were pooled and cleaned up using AMPure XP beads (Beckman-Coulter, A63881). Purity was validated using gel electrophoresis with a TBE gel. The purified products were spiked with 15% PhiX (Illumina, FC-110-3001) and sequenced using paired-end sequencing (read 1: 28 cycles, read 2: 66 cycles, i7 index: 8 cycles, i5 index: 8 cycles) on an Illumina NextSeq 2000 system (donors 1 and 2: P3 flow cell; donor 3: P2 flow cell).

## Data analysis

We analyzed 8504 CAR T cells at a single-cell level (Figure 1A), collected from 3 donors and stimulated once for acutely stimulated cells and stimulated repeatedly over 14 days for chronically stimulated cells, followed by selection for live cells and a surface-based marker NGFR indicating the expression of the CAR construct.

Data analysis was performed in python (version 3.11) using the pixelator^3^ (version 0.18), matplotlib, seaborn, scipy, and *scanpy* libraries.

### Data preprocessing

Sequencing results files were processed by *pixelator* to produce PXL files. These files contain preprocessed polarization and colocalization dataframes, as well as single-cell expression information in AnnData format.

### Calling cells

The PXL files contain a list of connections (edges) between neighboring pixels, which is used to construct a graph. Clustering this graph into components is performed using Leiden clustering within *pixelator*, and those components have varying numbers of edges. Components with more than 7000 edges were considered to be cells.

### Differential expression analysis

Differential expression analysis was performed using the *scanpy* (version 1.10.1) *rank_genes_groups_df* method (Table S2), and cutoffs of adjusted p-values less than 0.01 and log2 fold-change values greater than 1. These cutoffs were also applied to the determination of markers that were potentially confounded by trogocytosis, except the clusters were grouped into likely T cells and B cells prior to differential expression analysis.

### Dimensionality reduction and clustering

Dimensionality reduction was performed using the methods for principal component analysis (PCA) within the *scanpy* library. This was followed by louvain clustering using the *scanpy* library with resolution set to 1.

### Determining markers unique to NALM6 cells used to stimulate T cell response

The expression results are potentially confounded by trogocytosis of the markers on the NALM6 cells used to stimulate T cell response. To determine which markers may be affected by this process, we used a pilot dataset that did not employ FACS sorting of CAR T cell populations. Thus, this dataset included NALM6 cells, marked by the characteristic CD19 marker for B cells. We applied a similar pipeline as above but used the louvain clusters to distinguish likely T cells from B cells and applied differential expression between these populations. The cutoffs for the differential expression analysis are listed above, and the results are contained in Table S3.

For this pilot analysis, we performed standard live-cell counting with a hemocytometer and trypan blue staining. Samples 1, 2, 5, and 6 were >90% viable prior to fixation and freezing. Samples 3, 4, 7, and 8 where there was no tumor control had active killing of the target cells ongoing at the time of fixation, so the overall viability was around 50 % by live cell count due to dead cancer cells. Donor 1 (samples 1-4) was excluded from the analysis.

### Differential polarization analysis

Polarization testing removes control antibodies, subsets by donor and phenotype, and groups the polarization table by marker before dispatching each group to a parallel worker for scoring. The acute and chronic Moran’s-I z-score distributions for each marker are compared with a two-sided Mann–Whitney U test. The p-values of these tests were adjusted for multiple testing with the Bonferroni method, and markers with adjusted p-values < 0.001 were labelled significant for differential polarization and visualized in volcano-style summaries. This analysis also quantified each marker’s polarization variance and Gini coefficient across acute and chronic samples, using 2,000-permutation tests and Monte Carlo resampling against pooled background Moran’s-I values. The analysis was performed for CD8+ (Table S4), CD4+ (Table S5), and all CAR T cells (Table S6).

### Differential colocalization analysis

The colocalization workflow first drops self-pairs (included in the polarization analysis) and control antibodies. For each marker pair, the Pearson z-scores for acutely and chronically stimulated CAR T cells are compared with a two-sided Mann–Whitney U test. The p-values of these tests are adjusted for multiple testing using a Bonferroni correction. Pairs with z-score absolute values > 1 with adjusted p-values < 0.001 were marked as significant. This analysis also quantified each marker pair’s colocalization variance across acute and chronic samples by using permutation tests. This analysis was performed for CD8+ (Table S7), CD4+ (Table S8), and all CAR T cells (Table S9). Immune synapse interpretation layers were curated by zone assignments for anchor markers in the cSMAC, pSMAC, and dSMAC, and circos-style plots were generated to highlight partners meeting the significance thresholds.

### Variability analysis for polarization and colocalization

Both polarization and colocalization analyses analyzed the variability of these metrics across the 8504 single-cell measurements. This was performed by calculating the Gini coefficient across acute and chronic samples and performing permutation tests of this metric across 2,000 permutations. For each marker or marker pair, the observed Gini coefficient was computed from the original ordering, contrasted against 2,000 acute–chronic label permutations, and compared to 2,000 Monte Carlo resamples drawn from pooled background distributions that exclude the tested ordering. The resulting permutation test p-values for each marker, in polarization, or marker pair, for colocalization, were corrected for multiple testing with the Benjamini–Hochberg correction, and results with adjusted p-values below 0.01 were labeled to exhibit significant shifts in variability of polarization or colocalization over chronic stimulation of CAR T cells. The variability analysis results are reported for CD8+, CD4+, and all CAR T cells in Tables S10, S11, and S12, and the matrix used to construct Figure S5 is contained in Table S13. The variability analysis for colocalization did not reveal significant changes, and the results were omitted from this manuscript for brevity, although the code for performing this analysis is available in the provided repository.

## Supporting Figures

**Supporting Figure 1.**
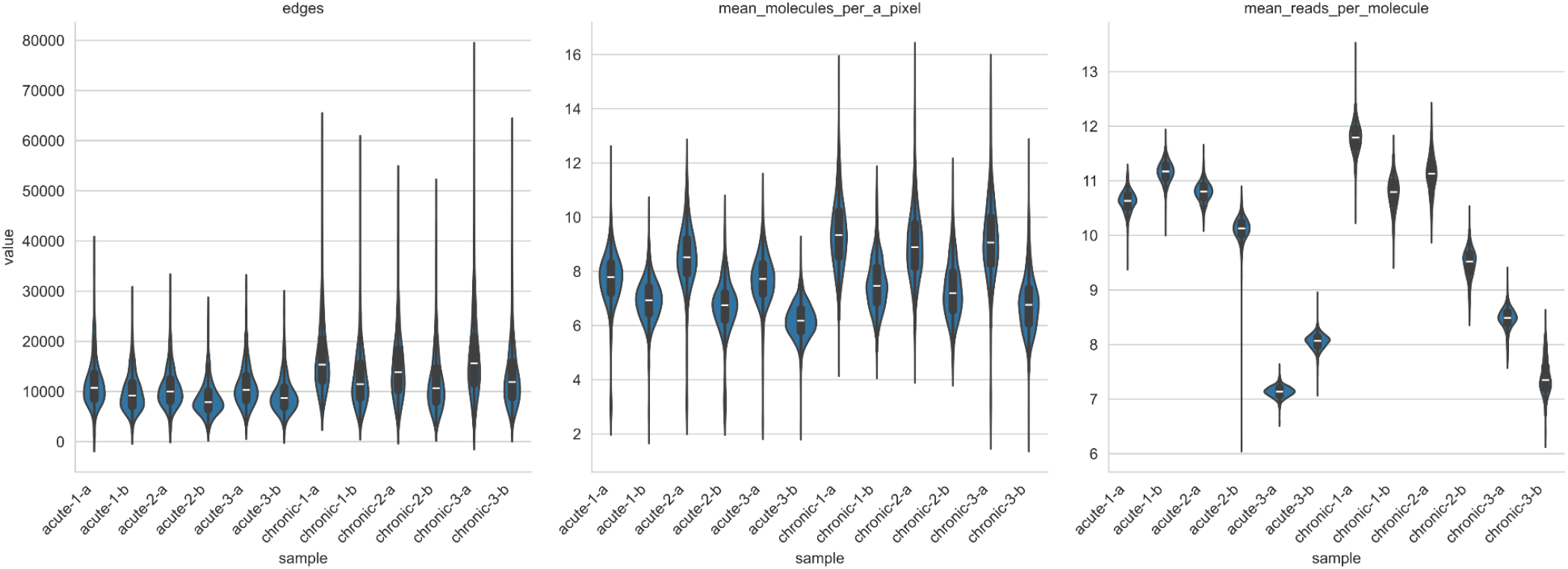
Quality control. A) Edges per cell, B) Mean molecules per pixel, C) Mean reads per molecule,

**Supporting Figure 2.**
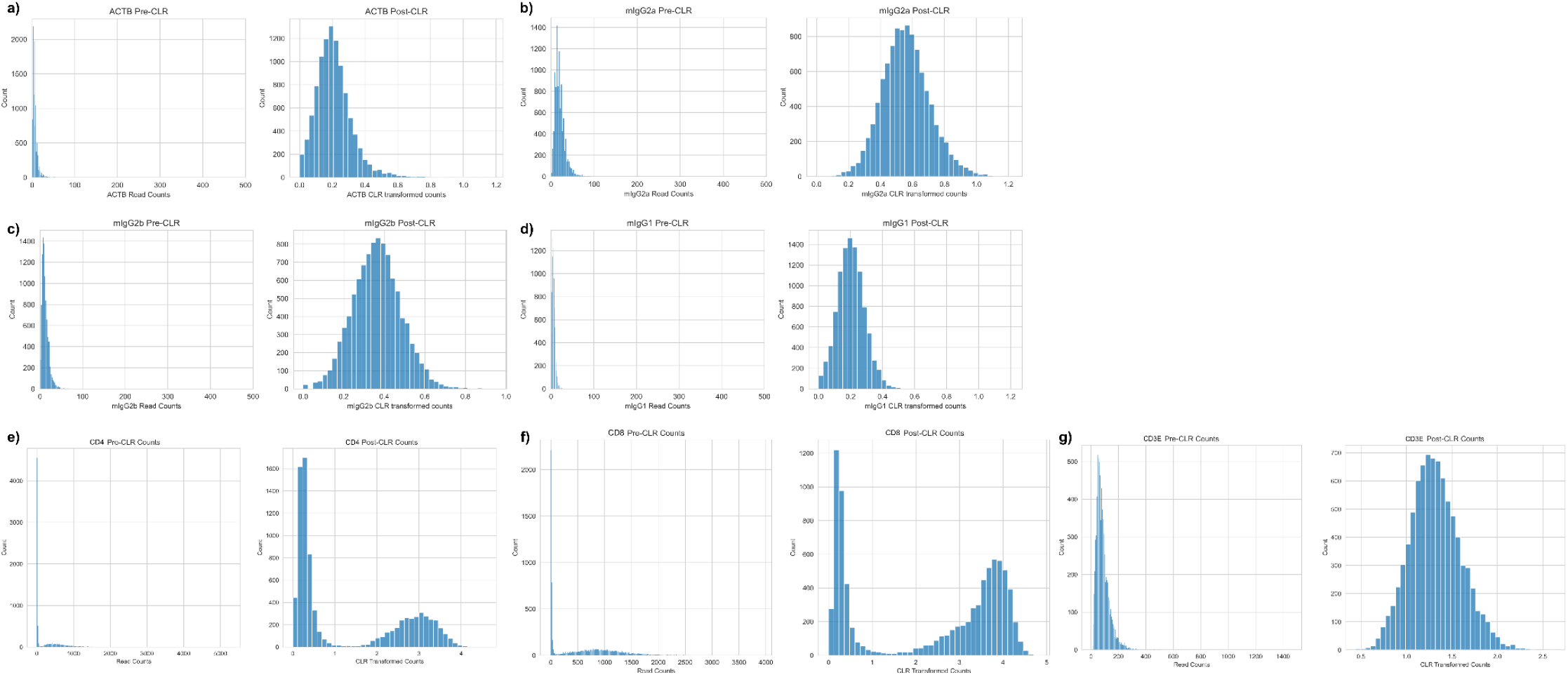
Normalization and comparison to controls. Intracellular control ACTB (a) and other controls mIG2a (b), mIG2b (c), and mIG1 (d) have low expression. Markers for CAR T cells CD4 (e) and CD8 (f) have bimodal populations representing two populations of these T cells, and the more ubiquitously expressed CD3E (g) is unimodal and highly expressed compared to the controls.

**Supplementary Figure 3.**
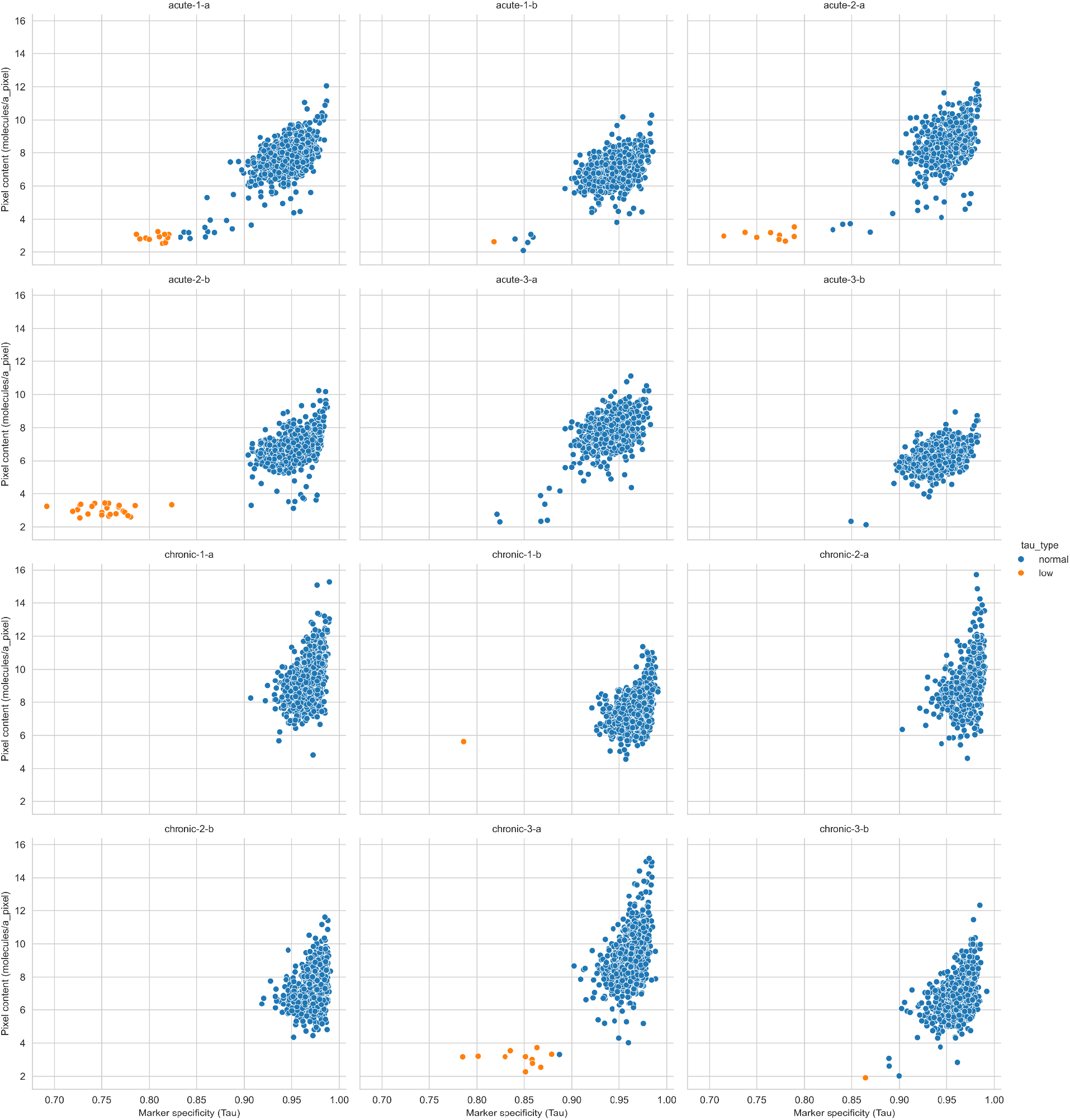
The tau metric for marker specificity is used to identify aggregates (orange) with low specificity that were filtered from the cells that were used for the analysis (blue) with higher marker specificity. Individual plots represent individual samples of acute stimulation (top two rows), and chronic stimulation (bottom two rows).

**Supplementary Figure 4.**
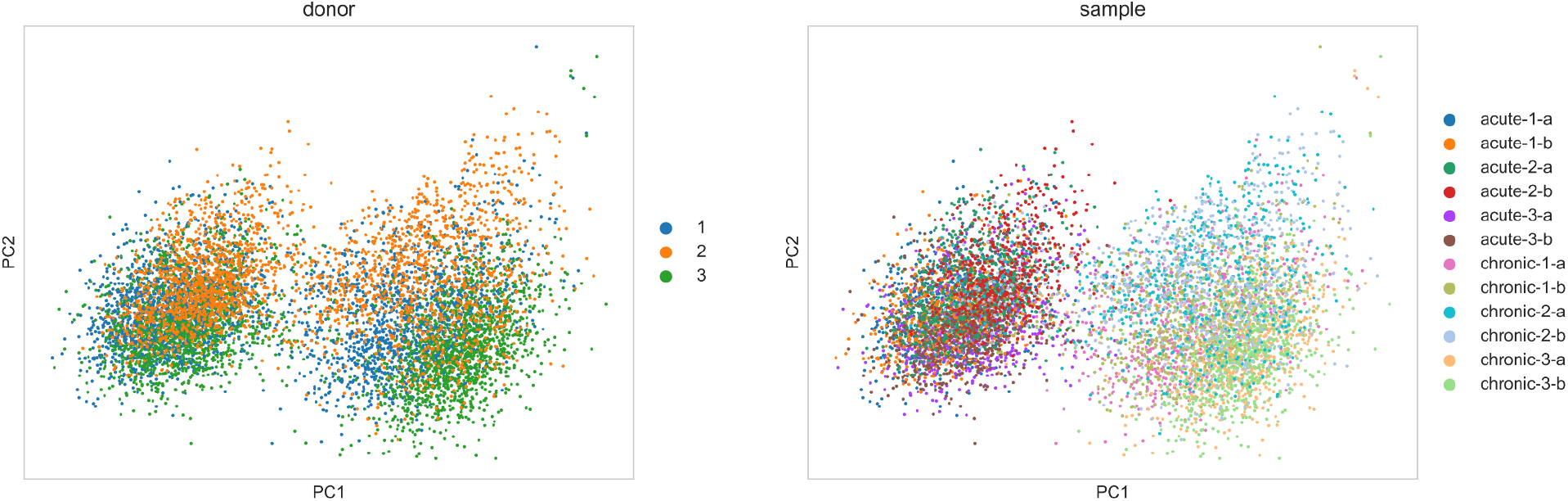
Dimensionality reduction plots using PCA showing the distribution of donors and samples.

**Supplementary Figure 5.**
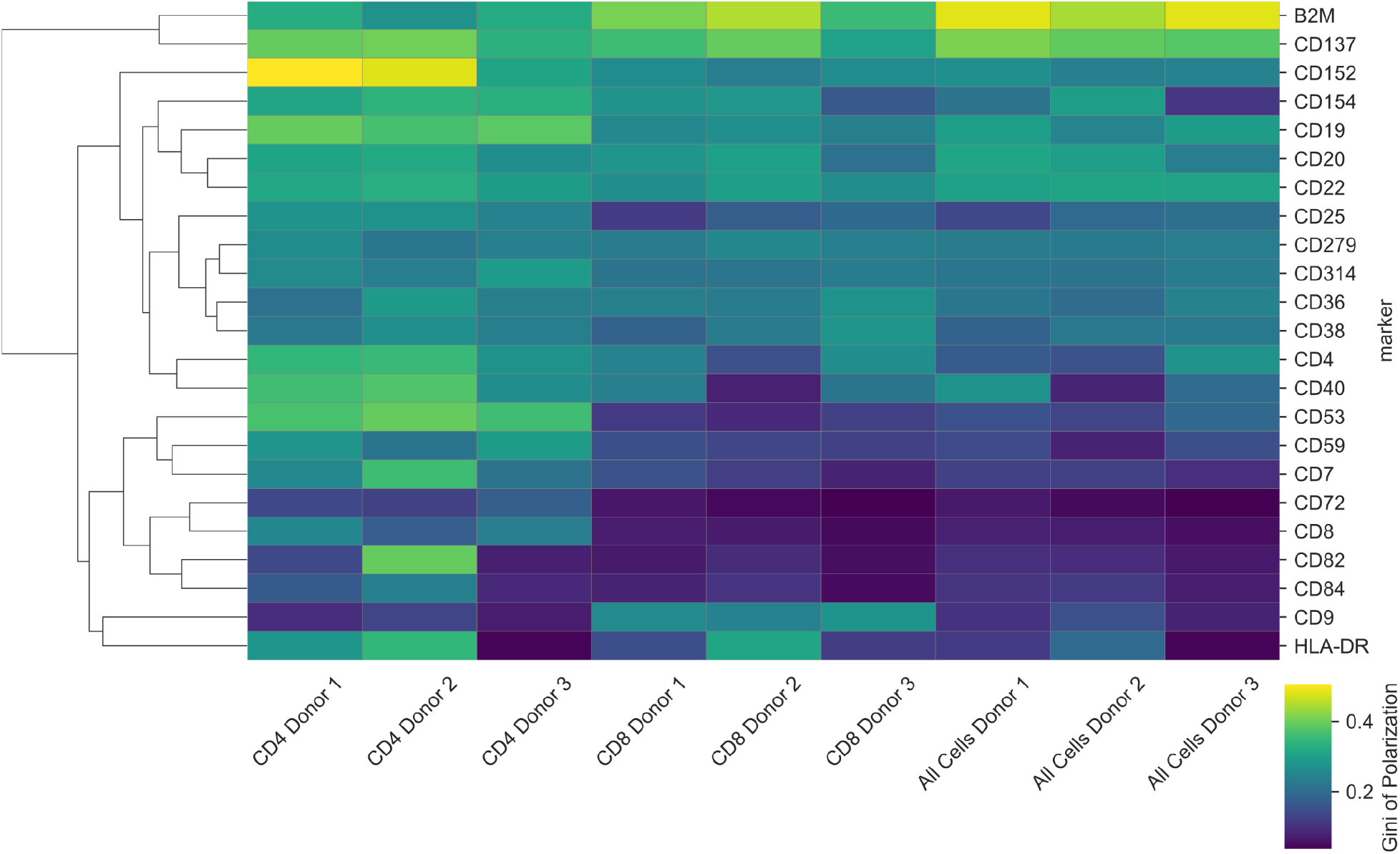
Clustermap of the single-cell variability (Gini index) of polarization for markers with significant changes in the variability of polarization for all CAR T cells in the study. This metric is relatively stable across the three donors.

## Acknowledgements

The authors thank the Pixelgen team for the help in sequencing, preprocessing the data, and conversations on data analysis. The Satpathy Lab was supported by the National Institutes of Health (NIH) U01CA260852, and A.C. was supported by Biohub’s donors, Priscilla Chan and Mark Zuckerberg.

## Author Contributions

A.C. and T.L. performed the data analysis. O.T.N. and T.L.R. prepared the samples. A.T.S. and E.L. coordinated funding.

## Competing Interests

T.L.R. is a founder of Arsenal Biosciences. A.T.S. is a founder of Immunai, Cartography Biosciences, Santa Ana Bio, and Arpelos Biosciences, an advisor to Wing Venture Capital, and receives research funding from Merck Research Laboratories, Allogene Therapeutics, and Astellas Pharma. E.L. is a founder of Genbio.ai and an advisor for Element Biosciences, Cartography Biosciences, Pfizer, Moleculent AB, and Pixelgen Technologies AB. The remaining authors declare no competing interests.

## Data and Materials Availability

The sequencing data is available on GEO SRA upon publication, along with the PXL files that were created using the data.

## Code Availability

The code available at https://github.com/CellProfiling/CAR-TCell-MPX-Analysis was used for this analysis and plot creation.

